# Modeling Vertical Migrations of Zooplankton Based on Maximizing Fitness

**DOI:** 10.1101/2021.01.29.428857

**Authors:** O. Kuzenkov, E. Ryabova, A. Garcia, A. Degtyarev

## Abstract

The purpose of the work is to calculate the evolutionarily stable strategy of zooplankton diel vertical migrations from known data of the environment using principles of evolutionary optimality and selection.

At the first stage of the research, the fitness function is identified using artificial neural network technologies. The training sample is formed based on empirical observations. It includes pairwise comparison results of the selective advantages of a certain set of species. Key parameters of each strategy are calculated: energy gain from ingested food, metabolic losses, energy costs on movement, population losses from predation and unfavorable living conditions. The problem of finding coefficients of the fitness function is reduced to a classification problem. The single-layer neural network is built to solve this problem. The use of this technology allows one to construct the fitness function in the form of a linear convolution of key parameters with identified coefficients.

At the second stage, an evolutionarily stable strategy of the zooplankton behavior is found by maximizing the identified fitness function. The maximization problem is solved using optimal control methods. A feature of this work is the use of piecewise linear approximations of environmental factors: the distribution of food and predator depending on the depth. As a result of the study, mathematical and software tools have been created for modeling and analyzing the hereditary behavior of living organisms in an aquatic ecosystem. Mathematical modeling of diel vertical migrations of zooplankton in Saanich Bay has been carried out.

## 1 Introduction

The phenomena of daily recurring vertical migrations of zooplankton were discovered more than two hundred years ago [1]. The study of the marine zooplankton’s behavior is of great importance due to zooplancton is a key link in the food chain. It plays a decisive role in the aquatic ecosystem; its diel migrations represent one of the most significant synchronous movements of biomass on earth. As a result, they affect carbon exchange and the climate of the planet [2–5]. In this regard, the problem of mathematical modeling of zooplankton’s diel vertical migrations is of great importance [6–12].

Currently, Darwin’s idea “survival of the fittest” is effectively used for modeling biological processes [13, 14]. It is possible to predict the results of evolution and to study the direction of changes in ecological systems comparing fitness of different biological species. Maximizing the fitness function provides the possibility to identify evolutionarily stable hereditary behavioral strategies (i.e. strategies that persist in the community against the appearance of possible mutations [15]). In particular, the use of the fitness concept for modeling diel migrations of zooplankton provides the opportunity to explain the quantitative characteristics of the behavior and its dependence on the age of an individual [16–18]. In this case, the main difficulty is the identification of the fitness function and its parameters.

There is a general approach to solving this problem based on studying the dynamics of a population distribution over the space of hereditary elements. This approach was proposed in [19] and was further developed in a series of works [20–22]. It was shown that on the set of hereditary elements it is possible to introduce a partial ranking order reflecting selective advantages by analyzing the long-term dynamics of the corresponding numbers of individuals [23]. The fitness function is introduced as a comparison function expressing the given ranking order. Then the problem of identifying the fitness function is reduced to expressing this function through the known hereditary features of elements.

In [24], the methodology for deriving the mathematical expression of the fitness function was developed for wide classes of population models, taking into account age heterogeneity. However, the parameters and coefficients of the model cannot quite often be measured empirically, and by themselves presuppose identification making the restoration of the fitness function much more difficult. Therefore, it seems interesting to construct the fitness function directly on the basis of the known population dynamics. In this case, the problem of restoring the fitness function is a special case of the well-known ranking problem [25]. For its solution, there is a wide arsenal of computer methods, in particular, machine learning methods (learning-to-rank) [26–32]. In [33, 34], the problem of ranking hereditary elements and identifying the corresponding fitness function was reduced to the problem of classification - dividing ordered pairs of elements into two classes: “the first element is better than the second” and “the second element is better than the first”.

In this work, this technique is used to identify the fitness function of diel vertical migrations of zooplankton. Parameters of the fitness function are identified on the basis of empirical observations. A feature of this work is the use of piecewise linear approximations of the distribution of food and predator depending on the depth of immersion. An evolutionarily stable behavior strategy is found by maximizing the identified fitness function using optimal control methods. As a result, mathematical modeling of diel vertical migrations of zooplankton in Saanich Bay is carried out.

## 2 Materials and methods

The present study is based on the following methodology for comparing the selective advantages of hereditary elements (behavior strategies) [24]. Let some compact metric space *V* of hereditary elements *v* be given. For example, such elements *v* can be continuous functions. Each element *v* at each moment of time *t* is assigned a certain number *ρ*(*v, t*) (indicator of presence), which numerically characterizes the presence of *v* in the community at time *t*. The indicator of presence satisfies the following requirements: it is zero when the element is not present in the community; it is strictly larger than zero when the element is presented to the community; the indicator is continuously dependent on time; its tendency to zero corresponds to the loss (extinction, disappearance) of this element in the community. This indicator can be the number, biomass of the subpopulation with a given hereditary element, the density of distribution of the population in the space of hereditary elements, etc.

Using the introduced indicator of presence, the selective advantages of various hereditary elements are compared with each other, namely, it is considered that the element *v* is better than the element *w* if

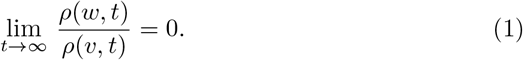

In the case when the presence indicator is uniformly above bounded (the community size is uniformly above bounded), the limit (1) means that the element *v* displaces the element *w* from the community over time. Thus, a partial order of selective advantages is given on the set *V*.

It is assumed that the introduced order can be expressed using the comparison functional *J*(*v*), that is, there is a functional that satisfies the condition *J*(*v*) > *J*(*w*) if and only if *v* is better than *w*. Then the functional *J* is a fitness function reflecting the selective advantages of hereditary elements.

If the change of the presence indicator in time is uniquely determined by a finite set *M*(*v*) = (*M*_1_(*v*),…, *M_n_*(*v*)) of key hereditary parameters (features) of the element *v*, then the functional *J* will be a function of these parameters: *J*(*v*) = *J*(*M*(*v*)). If this function is sufficiently smooth, then it is expedient to use Taylor’s expansions for its approximation. The simplest approximation is a linear convolution of key parameters

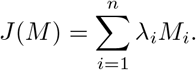

Here, the weights *λ_i_* reflect the impact of each key parameter on overall fitness. The problem of identifying the fitness function is reduced to determining the values of the convolution coefficients.

If it is known that the element *v* is better than the element *w* (from the analysis of the dynamics of the presenceindicator), then the in equality *J*(*M*(*v*)) > *J*(*M*(*w*)) should be fulfilled, respectively, the coefficients *λ_i_* should satisfy the inequality

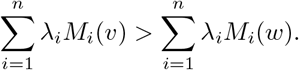

Knowing the results of comparing hereditary elements from a certain finite set, one can build a system of linear inequalities with respect to the convolution coefficients, which can be solved using linear programming methods [17].

Nevertheless, identification of these coefficients is also possible based on classification methods [33, 34]. Let us associate an ordered pair of elements (*v, w*) with a point *M*(*v*) – *M*(*w*), a pair (*w, v*) with a point *M*(*w*) – *M*(*v*) in the *n*-dimensional space of key parameters. Then the hyperplane

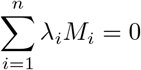

should separate these points from each other. A certain set of pairs of hereditary elements with known comparison results defines in a *n*-dimensional space two sets of points that must lie on opposite sides of this hyperplane. Thus, the problem of finding the convolution coefficients is reduced to finding the components of the normal to the separating hyperplane. This is the classification problem, for the solution of which there is a sufficient arsenal of well-proven methods [26]. For example, the separating hyperplane can be constructed using the Fisher determinant [35]. The classification problem is solved quite simply by the nearest neighbors method, but this method has limited application here, since it does not always allow one to find the coefficients of the separating hyperplane. One of the promising methods for solving this problem is the construction of a learning neural network [36, 37].

The formulated problem is also a special case of the pattern recognition problem [33, 34]. But in contrast to classical problems of this type, here it is not a simple assignment of an element to one of the two classes, but a comparison of elements according to the principle “better or worse”. Such a comparison is equivalent to recognizing the belonging of ordered pairs of elements “first, second” to one of two classes: “the first is better than the second” or “the first is worse than the second.”

As the experience of using various methods shows [34], the greatest effect can be obtained by using neural networks to solve the set problem. The neural networks technology provides greater flexibility of the algorithm with regard to expanding the training set, adding new experimental results of pair comparison. The use of neural networks provides a lower error rate compared to the nearest neighbors method.

It is known from the results of numerous studies that the main environmental factors affecting the behavior of zooplankton are: the degree of saturation of the water layer (with a vertical coordinate *x*) with food (phytoplankton) *E*_0_(*x*), metabolic costs *E*_2_(*x*) for maintaining viability in the water layer *x* (depends on the temperature of the layer), the number of predators (fish) *S_x_*(*x*) in the water layer *x*, the predator activity *S_t_*(*t*) depending on the time of day *t*, the presence of unfavorable factors *G*(*x*) in the water layer, such as temperature, hydrogen sulfide concentration, etc. [1, 6]. All of these factors are mathematically represented as functions of vertical coordinate or time.

Let us introduce a coordinate system so that *x* = 0 coincides with the water surface; *x* = −*D* is the level of the lethal hydrogen sulfide concentration (maximum immersion depth); *x* = −*C* is the level, below which there are neither predators feeding on zooplankton, nor phytoplankton, which feeds on zooplankton (*D*, *C* – positive constants, *C < D*). Let *t* be a time of day ranging from 0 to 1, with 0 being noon, 1/2 – midnight, 1 – next noon.

We take the following approximations of external factors:

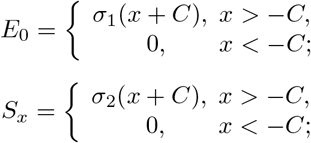

*S_t_* = cos 2*πt* + *∊* cos 6*πt* + 1; *S* = *S_x_ *·* S_t_*; *G* = *δ*(*x* + *D/*2)^2^; *E*_2_ = *σ*_3_(*x* + *D*).

In addition, it is assumed that the metabolic costs of zooplankton vertical migrations are proportional to the kinetic energy of movement, which in turn is proportional to the square of the speed: *E*_1_ = *ẋ*^2^.

On the one hand, the introduced functions *E*_0_, *E*_1_, *E*_2_, *S, G* represent a good approximation to the actually observed data, on the other hand, their relative simplicity allows us to investigate and solve the optimization problem analytically.

Figures 1-4 show the graphs of the functions *E*_0_(*x*), *G*(*x*), *S_x_*(*x*), *S_t_*(*t*) at *D* = 120, *C* = 50, *∊* = −0.013, *σ*_1_ = −0.018, *σ*_2_ = 1.8, *σ*_3_ = 1, (which corresponds to the data of empirical observations [1, 6, 18]).

**Fig. 1.**
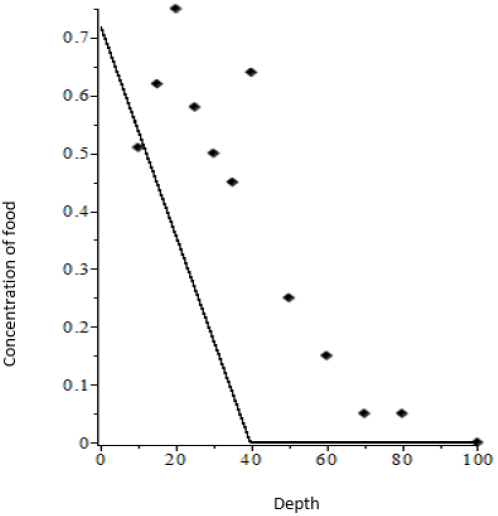
Amount of food (*E*_0_(*x*))

**Fig. 2.**
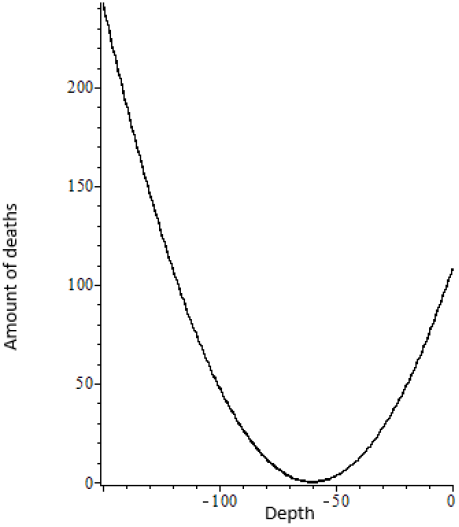
Additional mortality caused by approaching habitat boundaries (*G*(*x*))

**Fig. 3.**
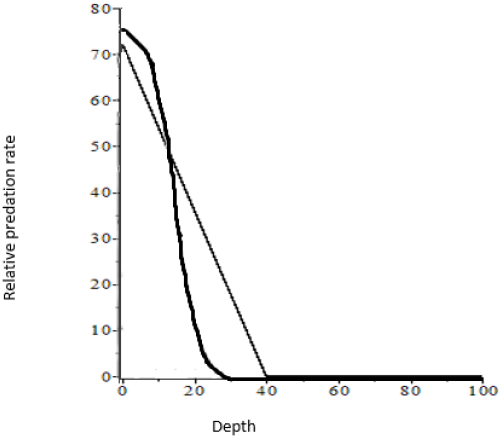
Mortality due to predation *S_x_*(*x*)

**Fig. 4.**
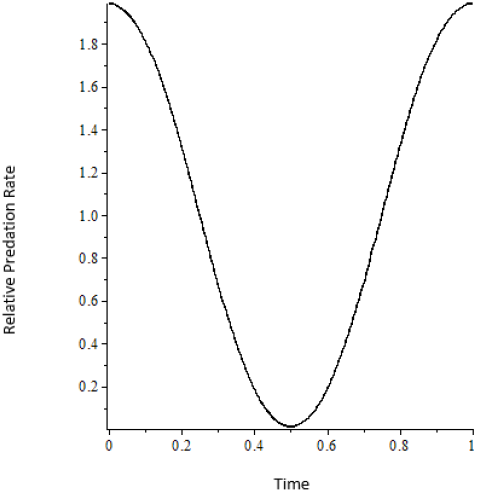
Number of attacks in time (*S_t_*(*t*))

## 3 Results

### 3.1 Fitness identification

The linear approximations of the fitness function were built using a neural network.

Let *x*(*t*) be the hereditary strategy of the zooplankton behavior, the depth of immersion depending on the time of day. It is obvious that the function *x*(*t*) must be continuous periodic with a period *T* = 1 (one day). This implies the condition x(0) = x(1). It is also assumed that this function is smooth.

It is possible to calculate the key parameters of the behavioral strategy *v* on the base of known functions of external factors

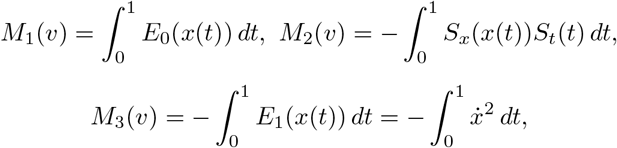

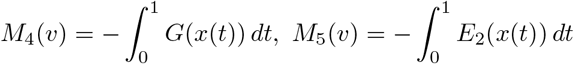

and the corresponding vector *M*(*v*) = (*M*_1_(*v*), *M*_2_(*v*), *M*_3_(*v*), *M*_4_(*v*), *M*_5_(*v*)).

It is assumed that the fitness function depends on these parameters linearly as follows

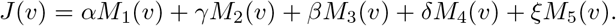

or

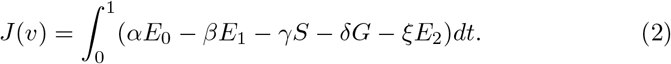

Weighting coefficients *α, γ, β, δ, ξ* determine the impact of each factor on overall fitness. The problem is to find the values of these coefficients.

To solve this problem, it is necessary to use information about known strategies of behavior. We can compare strategies *v* and *w* with each other, if we know the long-term dynamics of corresponding indicators *ρ*(*v, t*) and *ρ*(*w, t*). Then we can use the described above technology to estimate the coefficients *α, γ, β, δ, ξ* on the base of comparison results for a certain set of pairs.

To solve this problem, a single-layer neural network was built, which allows us to recognize pairs of hereditary strategies by their belonging to two classes - “the first strategy is better than the second” or “the second strategy is better than the first”.

This mathematically corresponds to constructing a hyperplane in a five-dimensional space separating two sets of points.

The coordinates of the normal of the constructed hyperplane correspond to the values of the required coefficients *α*, *γ*, *β*, *δ*, *ξ*.

For the computer solution using neural network technologies, the following standard free software was used: Scikit-learn machine learning library for the Python programming language, Pandas software library in Python for data processing and analysis.

The training sample was built taking into account the empirical results of observing the behavior of zooplankton ([18, 38]). It contains comparing results for 202 strategies or 2031 pairs.

The training sample was divided at a percentage of 70% for training by 30% for testing using the train test split module from the sklearn.model selection library. The quality of training was assessed using the Logloss metric. The learning error in this metric is 9.99e-16. The second check method was also used, using the cross val score function from the sklearn.model selection library. The recognition is performed with an accuracy of 96.3%.

Figure 5 shows a visualization of the solution to the corresponding classification problem.

**Fig. 5.**
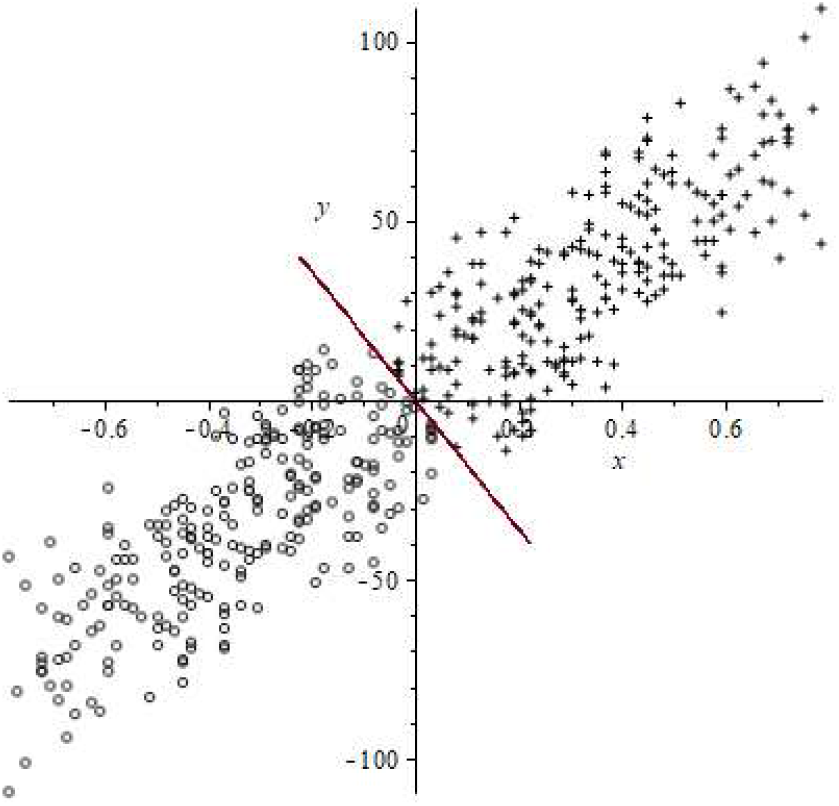
Solution of the linear classification problem for two classes of pairs of strategies.

Here the projections of the points of the training sample are shown. They correspond to different pairs of strategies onto the plane of two key parameters that have the meaning of food consumed per day – *M*_1_ and daily losses from predators – *M*_2_. The projections have coordinates (*M*_1_(*v*) − *M*_1_(*w*), *M*_2_(*v*) − *M*_2_(*w*)). The crosses mark the points corresponding to the pairs (*v, w*) for which *v* is better than *w*; the circles mark the points for which *v* is worse than *w*. The straight line corresponds to the intersection of the separating hyperplane and the plane of the parameters *M*_1_ and *M*_2_. The graph shows that the hyperplane accurately separates two classes of points from each other.

Found values of fitness coefficients are *α* = 344.444, *β* = 3.25·10^*−*5^, *γ* = 1.461, *δ* = 0.03, *ξ* = 2.24.

### 3.2 Optimization problem solution

The problem of constructing the evolutionarily stable strategy for zooplankton was solved as an optimal control problem [16, 22, 39–41] by maximizing the fitness function (2).

Let us introduce the notation

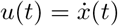

then the function *u* can be regarded as a control.

The conjugate system and transversality conditions have the following form [39]

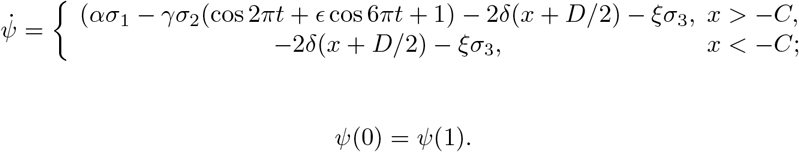

According to the minimum principle [39], the Hamilton function

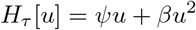

attains its minimum at the optimal control *u*(*τ*) for almost all times *τ*. Hence it follows that the optimal strategy *x*(*t*) of zooplankton behavior should satisfy the following conditions

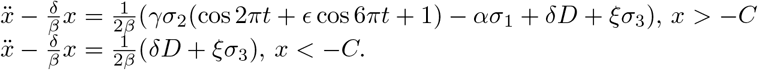

Note that the functional (2) is symmetric with respect to the replacement of the variable *t* by *τ* = 1 − *t*. Therefore, the solution in the interval 0 ≤ *t* ≤ 1 must be symmetric with respect to the time instant *t* = 1/2 and satisfy the condition *ẋ*(1) = −*ẋ*(0). Taking into account the periodicity of the solution, we conclude that *ẋ*(1) = *ẋ*(0) = 0.

Then the optimal solution *x*(*t*) is a continuous connection of functions

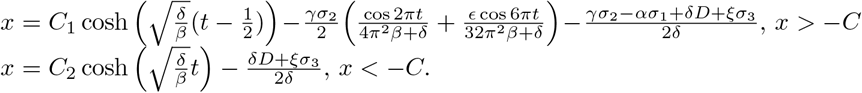

The constants *C*_1_ and *C*_2_ were calculated numerically to ensure a continuous connection. One can calculate the value of the constant *C*_2_ with a fixed arbitrary constant *C*_1_, at which two functions are continuously connected. Then one can choose such *C*_1_, at which functional (2) reaches its maximum. The standard Maple 17 package was used to solve the problem numerically.

Fig.6 shows the calculated trajectory of zooplankton movement in comparison with the empirically observed strategy of vertical movement of zooplankton in Saanich Bay [38]. Found constants are *C*_1_ = *−*0.003, *C*_2_ = 0.054.

**Fig. 6.**
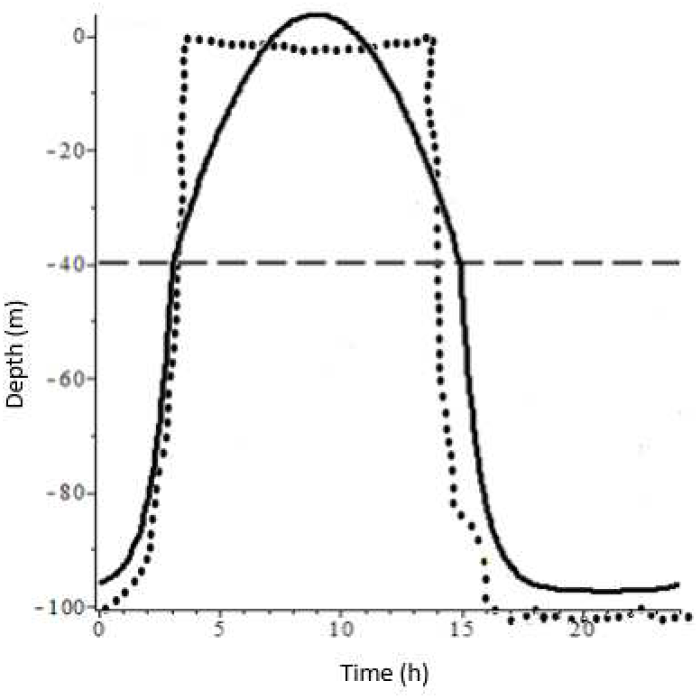
Comparison with experimental data obtained on 01.04.2010 from Saanich. The dotted line indicates the path most likely followed by zooplankton, and the continuous line is the line obtained by our model with *D* = 120, *C* = 50, *α* = 344.444, *β* = 3.25 *·* 10^*−*5^, *γ* = 1.461, *δ* = 0.03, *ξ* = 2.24, *∊* = *−*0.013, *σ*_1_ = 0.018, *σ*_2_ = 1.8, *σ*_3_ = 1 and constants *C*_1_ = *−*0.003, *C*_2_ = 0.054.

## 4 Summary

This study continues a series of works by the authors devoted to modeling the behavior of a zooplankton population using the principles of evolutionary optimality and selection. It is shown how artificial neural networks can be used to identify the fitness function of living organisms. The fitness function is built on the basis of pairwise comparison of the selective advantages of a certain set of species. The problem of finding the coefficients of the fitness function is reduced to the problem of classification. The parameters of the fitness function are identified on the basis of empirical observations.

Mathematical and software tools have been created for modeling and analyzing the hereditary behavior of living organisms in an aquatic ecosystem, determining their evolutionarily stable strategy and predicting changes in the system.

A feature of this work is the use of piecewise linear approximations of the distribution of food and predator depending on the depth of immersion. An evolutionarily stable strategy of zooplankton behavior is found by maximizing the identified fitness function by optimal control methods. As a result, mathematical modeling of diel vertical migrations of zooplankton in Saanich Bay is carried out.

It should be noted that results of work were implemented in the educational process of Lobachevsky State University of Nizhny Novgorod. The results are used within studying of the discipline “Mathematical modeling of selection processes” [42, 43]. They are used for the providing final qualification works of bachelors and masters. It provides the close connection of science and education and corresponds to the modern trends of the education modernization [44].

